# Distinct cognitive and structural correlates of pain extent and central sensitization symptoms in older women and men with chronic pain

**DOI:** 10.64898/2026.08.03.742487

**Authors:** Keiko Yamada, Hiroki Tabata, Kaito Takabayashi, Hitoshi Naito, Hideyoshi Kaga, Koji Kamagata, Yoshifumi Tamura

## Abstract

Chronic pain in later life may be accompanied by alterations in brain structure and cognition, but whether pain extent and central sensitization symptoms identify distinct brain-behavior patterns remains unclear. We examined associations of pain extent and central sensitization symptoms, assessed using the 9-item Central Sensitization Inventory (CSI-9), with regional gray matter volume and cognitive function in community-dwelling older adults. This cross-sectional study included 272 participants with chronic pain from the Bunkyo Health Study. Participants were classified as having single-site or multisite pain and by CSI-9 score as having lower or higher scores, with 12 or higher defining the higher group. Regional gray matter volume was quantified using 0.3-Tesla magnetic resonance imaging, and cognition was assessed using the Trail Making Test Part B (TMT-B), processing speed, and global and domain-specific measures. Pain extent and CSI-9 group interacted for TMT-B performance, with the longest completion time in participants with single-site pain and a higher CSI-9 score. No other cognitive outcome remained significant after correction for multiple testing. In categorical analyses, the higher CSI-9 group had smaller volumes in the right middle frontal gyrus, bilateral anterior cingulate cortex, right insula, right hippocampus, and bilateral amygdala, whereas pain extent and the interaction were not associated with regional volume. In a contextual comparison, only the single-site/higher CSI-9 group showed slower TMT-B performance than participants with no pain. Pain extent and central sensitization symptoms may represent partly distinct dimensions of chronic pain, although the small single-site/higher CSI-9 group and attenuation in several sensitivity analyses warrant caution.

**Significance Statement:** Chronic pain is often described by where it hurts, but location alone may miss important differences between patients. In older adults, pain extent and symptoms measured by the 9-item Central Sensitization Inventory captured partly different aspects of chronic pain. Participants with pain at one site and a higher CSI-9 score performed most slowly on a task requiring attention and mental flexibility, whereas differences in regional brain structure were related mainly to CSI-9 score rather than pain extent. These findings support a multidimensional approach to chronic pain and may inform future research on cognitive vulnerability and brain health across pain conditions.

**Graphical Abstract Text:** 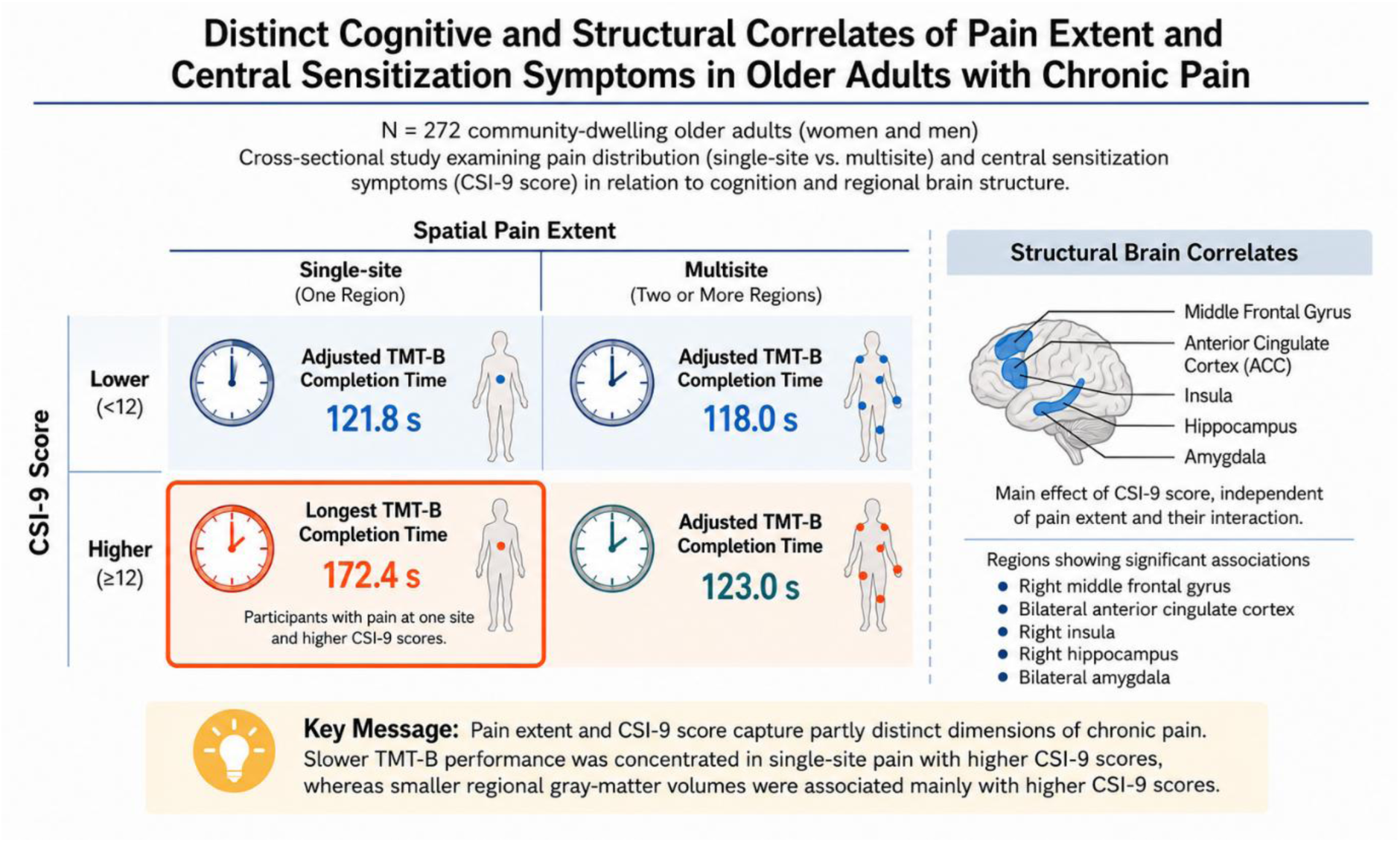

Among older adults with chronic pain, pain extent and CSI-9 score captured different aspects of vulnerability. Slower performance on a task requiring attention and cognitive flexibility was concentrated in those with single-site pain and higher CSI-9 scores, whereas regional brain-volume differences tracked CSI-9 category more broadly.

## 1. Introduction

Chronic pain is common in later life and contributes substantially to disability, reduced quality of life, and health-care burden (Cohen et al., 2021; Domenichiello & Ramsden, 2019). Contemporary mechanism-based frameworks distinguish nociceptive, neuropathic, and nociplastic pain mechanisms, which can coexist within the same individual. Nociplastic pain describes pain arising from altered nociception that is not fully explained by nociceptive or neuropathic mechanisms. Although central sensitization may contribute to nociplastic pain, the two concepts are not interchangeable (Kaplan et al., 2024; Kosek et al., 2021). The Central Sensitization Inventory is a patient-reported measure of symptoms commonly associated with central sensitization-related conditions; it does not directly quantify central nociceptive responsiveness. Its nine-item short form (CSI-9) provides a brief assessment of these symptoms (Mayer et al., 2012; Neblett et al., 2024; Nishigami et al., 2018).

Pain extent and CSI-9 score may capture related but partly distinct dimensions of the chronic pain phenotype. In community-based studies, higher CSI scores have been associated with a greater number of painful sites, greater pain severity, and multisite pain (Hoshino et al., 2022; Inoue et al., 2025). These associations do not imply that spatial pain extent and central sensitization symptoms are equivalent constructs. Accordingly, a higher CSI-9 score is not conceptually restricted to multisite pain, and multisite pain does not necessarily imply a higher CSI-9 score. Examining these dimensions jointly may identify clinically relevant heterogeneity that is obscured when chronic pain is classified only by diagnosis or by the number of painful sites (Henn et al., 2023; Kaplan et al., 2024).

Neuroimaging studies have linked chronic pain to structural differences across distributed brain systems involved in nociception, salience detection, affective processing, memory, attention, and cognitive control. However, recent evidence indicates that reported gray-matter differences are generally modest, spatially distributed, and heterogeneous across pain conditions and analytic methods, rather than reflecting a single pain-specific neural signature. Population-based work across different chronic pain conditions has also reported smaller gray-matter volumes in the anterior cingulate cortex, insula, and hippocampus (Neumann et al., 2023). Most structural imaging studies have compared individual pain diagnoses with pain-free controls, whereas fewer have examined whether pain extent and central sensitization symptoms make separable contributions to regional gray-matter volume. Such analyses are particularly relevant for frontal, cingulate, insular, hippocampal, and amygdalar regions involved in overlapping cognitive, affective, salience, and memory functions.

Cognitive function is also an important consideration in older adults with chronic pain. Longitudinal and large population-based studies have associated chronic pain, particularly pain at multiple sites, with poorer cognitive performance, cognitive decline, and smaller hippocampal volume (Rouch et al., 2021; Zhao et al., 2023). Selective-attention research further suggests that chronic pain can alter the allocation of attention toward pain-related information, although the magnitude and consistency of these effects vary across tasks and contexts (Abudoush et al., 2023). The Trail Making Test Part B is relevant because its performance depends on attentional control, set shifting, visual search, and processing speed. In older populations, cerebral small-vessel disease can also contribute to executive and attentional performance and to structural brain differences, making vascular imaging markers an important alternative explanation for observed pain-related associations (Jansma et al., 2024).

To address these gaps, we analyzed data from the Bunkyo Health Study (BHS), a community-based cohort of older adults in Japan (Someya et al., 2019). Among participants with chronic pain, we examined pain extent (single-site vs multisite) and CSI-9 score category (lower vs higher) as two factors. We tested their main and interactive associations with Trail Making Test Part B performance and prespecified regional gray-matter volumes, while treating other cognitive measures as secondary outcomes. We hypothesized that higher CSI-9 scores would be associated with slower Trail Making Test Part B performance and smaller regional gray-matter volumes and tested whether these associations differed between participants with single-site and multisite pain. Given that cerebral small-vessel disease can affect both cognition and brain structure in older adults, vascular brain pathology represents an important alternative explanation for any observed associations. We also sought to determine whether the identified chronic-pain phenotype differed from older adults reporting no current pain, thereby placing the findings in a broader clinical context.

## 2. Methods and Materials

### 2.1 Study Design and Participants

The cross-sectional study used data from the 5-year follow-up survey of the BHS, conducted between 2021 and 2024. The BHS is a prospective community-based cohort study of older adults residing in Bunkyo-ku Ward, Tokyo, Japan, and the detailed study design and recruitment procedures have been described previously (Someya et al., 2019). The study was approved by the Research Ethics Committee, Faculty of Medicine, Juntendo University in September 2015 (first approval no. 2015061, and the latest revised version no. M15-0057-M13), and all participants provided written informed consent.

Among 1,056 participants who participated in the on-site survey, 81 did not undergo brain magnetic resonance imaging (MRI), leaving 975 participants with MRI data. We subsequently excluded 88 participants with missing data on the pain questionnaire or face-to-face cognitive assessments, leaving 887 participants with valid MRI and basic assessment data. To reduce the potential influence of established dementia, participants were further excluded when they met both of the following operational criteria: a Mini-Mental State Examination (MMSE) score of ≤ 23 and a Montreal Cognitive Assessment-Japanese (MoCA-J) score of ≤ 22 (n = 17) (Fujiwara et al., 2010; Goldstein et al., 2014; Ideno et al., 2012).

After excluding a further 21 participants with missing data on pain duration or CSI-9 score, 849 participants (506 women and 343 men) were eligible for pain-status classification. Of these, 298 reported no current pain, 278 reported current pain lasting <3 months, and 273 reported chronic pain lasting ≥3 months. One participant with chronic pain endorsed none of the seven mapped pain regions and therefore could not be classified according to pain extent. The primary chronic-pain analysis consequently included 272 participants. The contextual TMT-B analysis combined these 272 participants (180 women and 92 men) with the 298 participants reporting no current pain, yielding an analytic sample of 570 participants (333 women and 237 men). The detailed enrollment process is shown in Figure 1.

**Figure 1.**
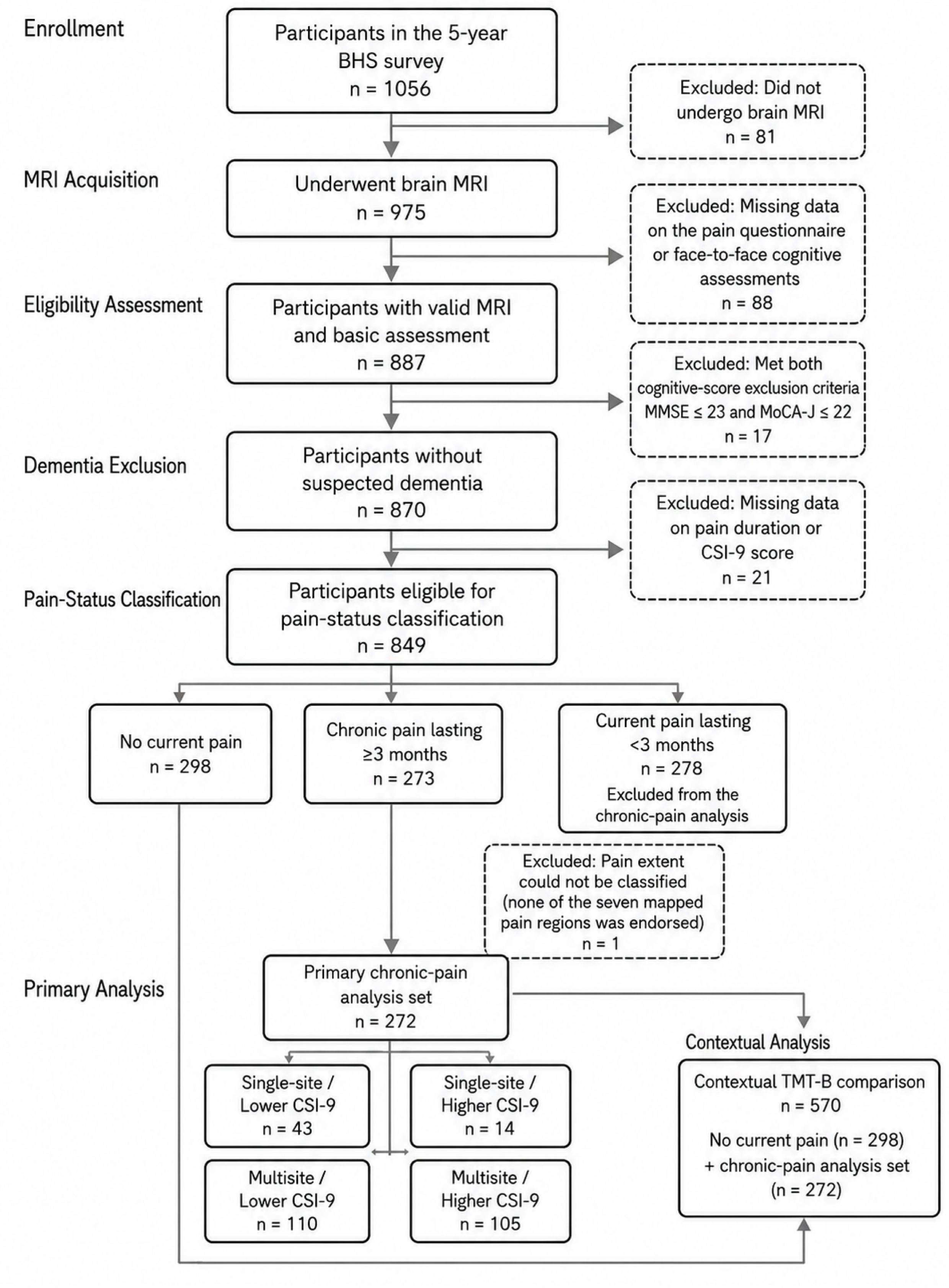
Participant flow and derivation of the analysis samples. The primary chronic-pain analysis included 272 participants with pain lasting ≥3 months and classifiable pain extent. Participants were grouped by pain extent and CSI-9 score. The contextual TMT-B analysis additionally included 298 participants with no current pain, yielding a total of 570 participants. Higher CSI-9 was defined as a CSI-9 score ≥12; single-site pain as exactly one of seven mapped pain regions; and multisite pain as two or more regions. **Abbreviations:** BHS, Bunkyo Health Study; CSI-9, 9-item Central Sensitization Inventory; MMSE, Mini-Mental State Examination; MoCA-J, Montreal Cognitive Assessment-Japanese version; MRI, magnetic resonance imaging; TMT-B, Trail Making Test Part B.

### 2.2 Pain Assessment and Phenotyping

Participants were classified according to the presence and duration of current pain. Chronic pain was defined as pain persisting for 3 months or longer (Treede et al., 2019). Participants who reported no current pain were classified as the no-current-pain group, whereas those with current pain lasting <3 months were excluded from the primary chronic-pain analysis.

Individual pain-location responses were mapped to seven body regions: the left upper extremity, right upper extremity, left lower extremity, right lower extremity, trunk, head/craniofacial/throat region, and visceral/pelvic region (Wolfe et al., 2016). The left-upper-extremity region included the left shoulder, left arm or upper arm, and left hand or fingers. Pain extent was classified as single-site pain when exactly one of the seven mapped regions was endorsed and as multisite pain when two or more regions were endorsed. A participant reporting chronic pain but endorsing none of the seven regions was considered to have unclassifiable pain extent and was excluded from the extent-based analyses. The distribution of pain regions is presented in Table S1.

The questionnaire also assessed average pain intensity and the degree to which pain affected general daily activity, each on a scale from 0 to 10, with higher scores indicating greater pain intensity or activity interference.

Symptoms associated with central sensitization were assessed using the 9-item Central Sensitization Inventory (CSI-9). Each item is rated from 0 to 4, yielding a total score ranging from 0 to 36, with higher scores indicating greater symptom severity (Nishigami et al., 2018). The primary categorical analysis defined a higher CSI-9 score as ≥12 and a lower CSI-9 score as <12. This prespecified cutoff corresponded to the upper quartile of the CSI-9 distribution in the full eligible sample rather than in the chronic-pain subgroup. Combining pain extent and CSI-9 score category yielded four groups: Single-site / Lower CSI-9, Single-site / Higher CSI-9, Multisite / Lower CSI-9, and Multisite / Higher CSI-9. The corresponding group sizes were 43, 14, 110, and 105 participants, respectively.

### 2.3 MRI Acquisition and Volumetric Processing

All participants underwent whole-brain MR imaging using a 0.3-T clinical MR scanner (AIRIS Vento, Hitachi, Tokyo, Japan). The following sequences were acquired: three-dimensional T1-weighted imaging using a gradient-echo inversion-recovery sequence (repetition time [TR] = 25 ms; echo time [TE] = 5.8 ms; inversion time [TI] = 600 ms; flip angle = 12°; number of excitations = 1; field of view = 200 × 250 × 250 mm³; voxel size = 0.98 × 0.98 × 2 mm³; slice orientation = sagittal; total scan time = 601 s) and fluid-attenuated inversion recovery (FLAIR) imaging (TR = 11,000 ms; TE = 100 ms; TI = 2000 ms; slice thickness = 5 mm) (Someya et al., 2019).

Although a lower magnetic field strength was used, a validation study by Murata et al. (2022) reported high agreement with good-to-excellent intraclass correlation coefficients across most brain regions between brain-volume measurements obtained using this 0.3-T scanner and those obtained using a standard 3-T scanner (MAGNETOM Prisma; Siemens Healthcare, Erlangen, Germany) in the same individuals, confirming the validity of this method for brain volumetry (Murata et al., 2022).

Image preprocessing and analysis were performed using voxel-based morphometry implemented in Statistical Parametric Mapping 12 (SPM12; RRID:SCR_007037) running under MATLAB R2022b (The MathWorks, Inc., Natick, MA, USA; RRID:SCR_001622). Voxel-based morphometry and DARTEL registration were performed as described previously (John Ashburner, 2007; John Ashburner & Friston, 2000). In the first step, T1-weighted images were segmented into six tissue classes: gray matter, white matter, cerebrospinal fluid, bone, soft tissue, and air. The segmented images were then resampled to 1.5-mm isotropic voxels and aligned across subjects using diffeomorphic anatomical registration through exponentiated Lie algebra (DARTEL). After DARTEL, the images were normalized to Montreal Neurological Institute (MNI) templates, with signal intensities modulated to preserve tissue volume. Finally, the images were smoothed with an 8-mm full-width-at-half-maximum Gaussian kernel. Spatial smoothing was performed to compensate for anatomical variability that was not fully accounted for by spatial normalization and to improve the signal-to-noise ratio (John Ashburner, 2007; John Ashburner & Friston, 2000; Murata et al., 2022).

Regional gray matter volumes were calculated using the Automated Anatomical Labeling (AAL) atlas (Tzourio-Mazoyer et al., 2002). The atlas was interpolated to 1.5-mm isotropic voxels to match the resolution of the modulated, normalized gray matter images, and regional gray matter volumes were obtained by summing the voxel values within each region of interest. We defined 10 regions of interest based on prior chronic-pain neuroimaging literature and their relevance to cognitive, affective, salience, and memory processes: the bilateral middle frontal gyrus, anterior cingulate cortex (ACC), insula, hippocampus, and amygdala. Regional volumes were expressed in cubic centimeters. Total intracranial volume was calculated as the sum of gray-matter, white-matter, and cerebrospinal-fluid volumes.

### 2.4 Cerebral Small-Vessel-Disease Markers

White matter hyperintensities (WMH) were visually graded on FLAIR images using the Fazekas scale (0–3) for periventricular and deep WMH (Fazekas et al., 1987). Lacunes of presumed vascular origin were identified according to the Standards for Reporting Vascular Changes on Neuroimaging criteria (round or ovoid, subcortical, cerebrospinal fluid-like cavities 3–15 mm in diameter, typically with a surrounding rim on FLAIR images). WMH/lacune ratings were performed by trained readers blinded to pain phenotype, with disagreements resolved by consensus (Wardlaw et al., 2013).

Periventricular hyperintensity grade, deep/subcortical white-matter hyperintensity grade, and lacune count were included as additional covariates in prespecified sensitivity analyses.

### 2.5 Cognitive Assessment

Cognitive assessments were administered face to face by trained psychologists. The primary outcome was the Trail Making Test Part B (TMT-B) completion time, expressed in seconds, with longer completion times indicating slower performance. TMT-B requires visual search, sustained attention, sequencing, and set shifting and was therefore selected as the primary measure of executive and attentional control.

Secondary cognitive outcomes were the TMT-A completion time, MMSE total score, MoCA-J total score, and the MoCA-J attention, visuospatial/executive, and abstraction domain scores. The Japanese versions of the MMSE and MoCA have been validated in older Japanese adults (Fujiwara et al., 2010; Ideno et al., 2012).

Because the maximum score is 30 for both the MMSE and MoCA-J, possible ceiling effects were assessed by calculating the proportion of participants attaining the maximum score within each of the four chronic-pain groups.

### 2.6 Statistical Analysis

Participant characteristics were summarized for the total chronic-pain sample and for each of the four pain-extent/CSI-9 groups. Continuous variables are presented as means and standard deviations, and categorical variables as counts and percentages. No missing values were imputed; analyses were based on participants with complete data for the relevant variables.

#### 2.6.1 Primary TMT-B analysis

The primary analysis used a 2 × 2 factorial analysis of covariance to examine the main effects of pain extent (single-site vs multisite), CSI-9 score category (lower vs higher), and their interaction on TMT-B completion time. Type III sums of squares were used because of the unequal group sizes. Models were adjusted for age, sex, educational attainment, Geriatric Depression Scale-15 score, history of cerebral infarction, history of cerebral hemorrhage, and current sleep-medication use. Educational attainment was entered using the categories 8–9, 10–12, 13–14, 15–16, and ≥17 years.

Adjusted marginal means and 95% confidence intervals were estimated for each of the four groups. Four prespecified simple comparisons were performed using Tukey–Kramer adjustment: higher versus lower CSI-9 within single-site pain, higher versus lower CSI-9 within multisite pain, single-site versus multisite pain within the higher CSI-9 category, and single-site versus multisite pain within the lower CSI-9 category. Partial eta-squared was reported as the effect-size measure.

#### 2.6.2 Secondary cognitive outcomes

The same factorial models and covariates were applied separately to TMT-A, MMSE total score, MoCA-J total score, and the MoCA-J attention, visuospatial/executive, and abstraction scores. To account for multiple testing, Benjamini–Hochberg false-discovery-rate correction was applied separately across the six secondary outcomes for each factorial effect. Results with an FDR-adjusted p value, denoted by *q*, <0.05 were considered statistically significant (Benjamini & Hochberg, 1995).

#### 2.6.3 Regional gray-matter volume analyses

Separate 2 × 2 factorial analyses of covariance were fitted for each of the 10 regional gray-matter volumes. These models included the same covariates as the cognitive models, with additional adjustment for total intracranial volume. Total intracranial volume was entered as a covariate rather than using proportional volume normalization (Wang et al., 2024).

For each of the three factorial effects—pain extent, CSI-9 score category, and pain extent × CSI-9 interaction, the Benjamini–Hochberg procedure was applied separately across the 10 regions of interest. An FDR-adjusted *q* value <0.05 was considered statistically significant. Partial eta-squared was reported for each effect.

#### 2.6.4 Contextual comparison with participants reporting no current pain

To place the primary chronic-pain findings in context, a separate analysis included the 298 participants reporting no current pain and the four chronic-pain groups, yielding five groups and a total sample of 570 participants. Participants with current pain lasting <3 months were not included. TMT-B completion time was compared across the five groups using analysis of covariance with the same covariates as the primary cognitive model. Dunnett-adjusted comparisons were used to compare each chronic-pain group with the no-current-pain reference group.

#### 2.6.5 Sensitivity and robustness analyses

For TMT-B, the robustness of the primary interaction was evaluated using the following alternative specifications: CSI-9 score modeled continuously; continuous painful-region count and continuous CSI-9 score with their interaction; a higher CSI-9 threshold of ≥16; multisite pain defined as ≥3 painful regions; additional adjustment for periventricular hyperintensity grade, deep/subcortical white-matter hyperintensity grade, and lacune count; log-transformed TMT-B completion time; TMT-B minus TMT-A; and the TMT-B/TMT-A ratio. A generalized estimating-equation model with empirical standard errors was also fitted using the same factorial terms and covariates.

Because the single-site/higher CSI-9 group included only 14 participants, a leave-one-out analysis was performed by repeating the primary TMT-B model 14 times, omitting one participant from this group at each iteration.

For regional gray-matter volumes, sensitivity analyses modeled CSI-9 score continuously, used the alternative CSI-9 cutoff of ≥16, defined multisite pain as ≥3 regions, and additionally adjusted for the three cerebral small-vessel-disease markers. False-discovery-rate correction was applied across the 10 regions separately for each factorial effect within each sensitivity analysis. All statistical tests were two-sided, with p<0.05 considered statistically significant before multiple-testing correction. Analyses were conducted using SAS version 9.4 (SAS Institute Inc., Cary, NC, USA, RRID:SCR_008567).

## 3. Results

### 3.1 Participant Characteristics

A total of 849 participants (mean age 76.9, 5.0 [standard deviation, SD] years; 59.6% female) were included in the analysis. Chronic pain was present in 273 participants (32.2%). One participant with chronic pain endorsed none of the seven mapped pain regions and was excluded because pain extent could not be classified. The primary chronic-pain analysis therefore included 272 participants. Their mean age was 77.42 years (SD, 4.99), and 180 participants (66.2%) were women.

On the basis of the phenotype classification, participants were classified into the single-site/lower CSI-9 (n=43), single-site/higher CSI-9 (n=14), multisite/lower CSI-9 (n=110), and multisite/higher CSI-9 (n=105) groups. In the total sample, the mean CSI-9 score was 11.01 (SD, 5.12), and the mean number of painful regions was 2.82 (SD, 1.41). Descriptively, the higher CSI-9 groups reported greater pain intensity, greater interference with general daily activity, and higher Geriatric Depression Scale-15 scores than the corresponding lower CSI-9 groups. Participant characteristics are presented in Table 1, and the distribution of the seven painful regions is shown in Table S1.

**Table 1.** Characteristics of participants with chronic pain according to pain extent and CSI-9 score.

| Characteristic | Total<br>(n=272) | Single-site /<br>Lower CSI-9<br>(n=43) | Single-site /<br>Higher CSI-9<br>(n=14) | Multisite /<br>Lower CSI-9<br>(n=110) | Multisite /<br>Higher CSI-9<br>(n=105) |
| --- | --- | --- | --- | --- | --- |
| **Age, years, mean (SD)** | 77.42 (4.99) | 76.00 (4.47) | 79.43 (6.35) | 77.64 (4.70) | 77.50 (5.21) |
| **Female sex, n (%)** | 180 (66.2) | 24 (55.8) | 11 (78.6) | 74 (67.3) | 71 (67.6) |
| **Educational attainment, n (%)** |  |  |  |  |  |
| 8–9 years | 12 (4.4) | 1 (2.3) | 0 (0.0) | 7 (6.4) | 4 (3.8) |
| 10–12 years | 112 (41.2) | 16 (37.2) | 11 (78.6) | 43 (39.1) | 42 (40.0) |
| 13–14 years | 47 (17.3) | 4 (9.3) | 1 (7.1) | 16 (14.5) | 26 (24.8) |
| 15–16 years | 95 (34.9) | 21 (48.8) | 2 (14.3) | 42 (38.2) | 30 (28.6) |
| ≥17 years | 6 (2.2) | 1 (2.3) | 0 (0.0) | 2 (1.8) | 3 (2.9) |
| **Pain-related characteristics, mean (SD)** |  |  |  |  |  |
| CSI-9 score (0–36) | 11.01 (5.12) | 5.95 (2.79) | 15.57 (3.16) | 7.92 (2.53) | 15.71 (3.37) |
| Number of painful regions | 2.82 (1.41) | 1.00 (0.00) | 1.00 (0.00) | 2.94 (1.00) | 3.68 (1.24) |
| Average pain intensity (0–10) | 3.83 (2.02) | 2.70 (1.96) | 4.93 (1.59) | 3.36 (1.85) | 4.64 (1.89) |
| General daily activity score (0–10) | 2.80 (2.18) | 1.67 (2.30) | 3.64 (1.65) | 2.26 (1.91) | 3.70 (2.08) |
| **Cognitive measures, mean (SD)** |  |  |  |  |  |
| TMT-A, sec | 44.75 (15.59) | 39.88 (10.71) | 47.64 (17.59) | 46.16 (17.30) | 44.87 (14.87) |
| TMT-B, sec | 119.88 (55.11) | 109.30 (52.29) | 183.21 (105.26) | 113.88 (42.99) | 122.04 (53.46) |
| MMSE total score (0–30) | 27.90 (1.62) | 27.67 (1.46) | 27.71 (1.86) | 28.16 (1.63) | 27.75 (1.63) |
| MoCA-J total score (0–30) | 25.94 (2.57) | 25.74 (2.41) | 25.29 (2.76) | 25.77 (2.53) | 26.29 (2.64) |
| **MRI and clinical covariates** |  |  |  |  |  |
| GDS-15 score (0–15), mean (SD) | 2.93 (2.47) | 1.74 (1.83) | 3.71 (2.20) | 2.21 (2.02) | 4.08 (2.67) |
| Total intracranial volume, L, mean (SD) | 1.40 (0.15) | 1.42 (0.16) | 1.41 (0.38) | 1.39 (0.11) | 1.39 (0.12) |
| History of cerebral infarction, n (%) | 7 (2.6) | 0 (0.0) | 0 (0.0) | 4 (3.6) | 3 (2.9) |
| History of cerebral hemorrhage, n (%) | 3 (1.1) | 0 (0.0) | 0 (0.0) | 3 (2.7) | 0 (0.0) |
| Current sleep-medication use, n (%) | 44 (16.2) | 3 (7.0) | 2 (14.3) | 15 (13.6) | 24 (22.9) |

**Abbreviation:** CSI-9, 9-item Central Sensitization Inventory; GDS-15, 15-item Geriatric Depression Scale; MMSE, Mini-Mental State Examination; MoCA-J, Japanese version of the Montreal Cognitive Assessment; SD, standard deviation; TMT-A, Trail Making Test, part A; TMT-B, Trail Making Test, part B.
**Note.** Higher CSI-9 was defined as $\geq 12$ . Single-site pain was defined as exactly one of seven mapped pain regions; multisite pain was defined as $\geq 2$ regions. n=272

### 3.2 Primary TMT-B Analysis

In the primary factorial analysis of covariance, TMT-B completion time was associated with pain extent (*F*=10.03, *P*=0.0017, partial η²=0.0374), CSI-9 group (F=9.92, p=0.0018, partial η²=0.0370), and the pain extent × CSI-9 group interaction (*F* =7.34, *P* =0.0072, partial η²=0.0276).

The covariate-adjusted mean TMT-B completion times were 121.80 seconds (95% CI, 103.95– 139.64) in the single-site/lower CSI-9 group, 172.41 seconds (95% CI, 143.98–200.83) in the single-site/higher CSI-9 group, 117.97 seconds (95% CI, 104.78–131.16) in the multisite/lower CSI-9 group, and 123.04 seconds (95% CI, 109.71–136.37) in the multisite/higher CSI-9 group.

Within participants with single-site pain, the higher CSI-9 group had a longer adjusted TMT-B completion time than the lower CSI-9 group (mean difference, 50.61 seconds; simultaneous 95% CI, 10.17–91.05; Tukey–Kramer *P*=0.0074). The corresponding difference within participants with multisite pain was small and not statistically significant (mean difference, 5.07 seconds; 95% CI, −13.76 to 23.90; *P*=0.8985). Among participants with a higher CSI-9 score, those with single-site pain had a longer completion time than those with multisite pain (mean difference, 49.37 seconds; 95% CI, 12.65–86.08; *P*=0.0033), whereas no difference according to pain extent was observed in the lower CSI-9 category (mean difference, 3.82 seconds; 95% CI, −19.42 to 27.06; *P*=0.9741). Thus, the association between a higher CSI-9 score and slower TMT-B performance was more pronounced among participants with single-site pain (Table 2).

**Table 2.** Primary analysis of Trail Making Test Part B performance according to pain extent and CSI-9 score.

| Type III tests |  |  |  |  |
| --- | --- | --- | --- | --- |
| Effect | df | F | P | Partial $\eta^2$ |
| Pain extent | 1 | 10.03 | 0.0017 | 0.0374 |
| CSI-9 group | 1 | 9.92 | 0.0018 | 0.037 |
| Pain extent × CSI-9 group | 1 | 7.34 | 0.0072 | 0.0276 |
| Covariate-adjusted TMT-B means |  |  |  |  |
| Group | n | Adjusted mean, sec | 95% CI |  |
| Single-site / Lower CSI-9 | 43 | 121.80 | 103.95 to 139.64 |  |
| Single-site / Higher CSI-9 | 14 | 172.41 | 143.98 to 200.83 |  |
| Multisite / Lower CSI-9 | 110 | 117.97 | 104.78 to 131.16 |  |
| Multisite / Higher CSI-9 | 105 | 123.04 | 109.71 to 136.37 |  |
| Prespecified simple comparisons |  |  |  |  |
| Comparison | Mean difference, sec | Simultaneous 95% CI | Tukey–Kramer P |  |
| Higher vs lower CSI-9 within single-site pain | 50.61 | 10.17 to 91.05 | 0.0074 |  |
| Higher vs lower CSI-9 within multisite pain | 5.07 | -13.76 to 23.90 | 0.8985 |  |
| Single-site vs multisite within higher CSI-9 | 49.37 | 12.65 to 86.08 | 0.0033 |  |
| Single-site vs multisite within lower CSI-9 | 3.82 | -19.42 to 27.06 | 0.9741 |  |

**Abbreviation:** CI, confidence interval; CSI-9, 9-item Central Sensitization Inventory.
**Note.** Models were adjusted for age, sex, education, 15-item Geriatric Depression Scale score, history of cerebral infarction, history of cerebral hemorrhage, and sleep-medication use. Positive differences indicate slower Trail Making Test Part B performance in the first listed group.

### 3.3 Secondary Cognitive Outcomes

None of the factorial effects for TMT-A, MMSE total score, MoCA-J total score, MoCA-J attention, MoCA-J visuospatial/executive function, or MoCA-J abstraction remained statistically significant after false-discovery-rate correction. The pain extent × CSI-9 group interaction for the MoCA-J visuospatial/executive score was significant before correction (*F*=6.66, *P*=0.0104, partial η²=0.0252) but did not meet the FDR-corrected threshold (*q*=0.0625). The other nominal and FDR-adjusted results are presented in Table S2.

### 3.4 Regional Gray Matter Volumes

Pain extent was not associated with regional gray-matter volume in any of the 10 prespecified regions after FDR correction. Similarly, no pain extent × CSI-9 group interaction remained significant after correction.

In contrast, the main effect of CSI-9 group was significant after FDR correction in seven regions: the right middle frontal gyrus (*F*=5.68, *q*=0.0356), left anterior cingulate cortex (*F* =5.36, *q*=0.0356), right anterior cingulate cortex (*F*=6.62, *q*=0.0356), right insula (*F*=5.57, *q*=0.0356), right hippocampus (*F*=5.78, q=0.0356), left amygdala (*F*=4.77, *q*=0.0427), and right amygdala (*F*=6.73, *q*=0.0356). Participants in the higher CSI-9 group had smaller adjusted gray-matter volumes in each of these regions. The corresponding effect sizes were small, with partial η² values ranging from 0.0182 to 0.0255. Full factorial results and adjusted regional means are presented in Table 3 and Table S4, respectively.

**Table 3.**
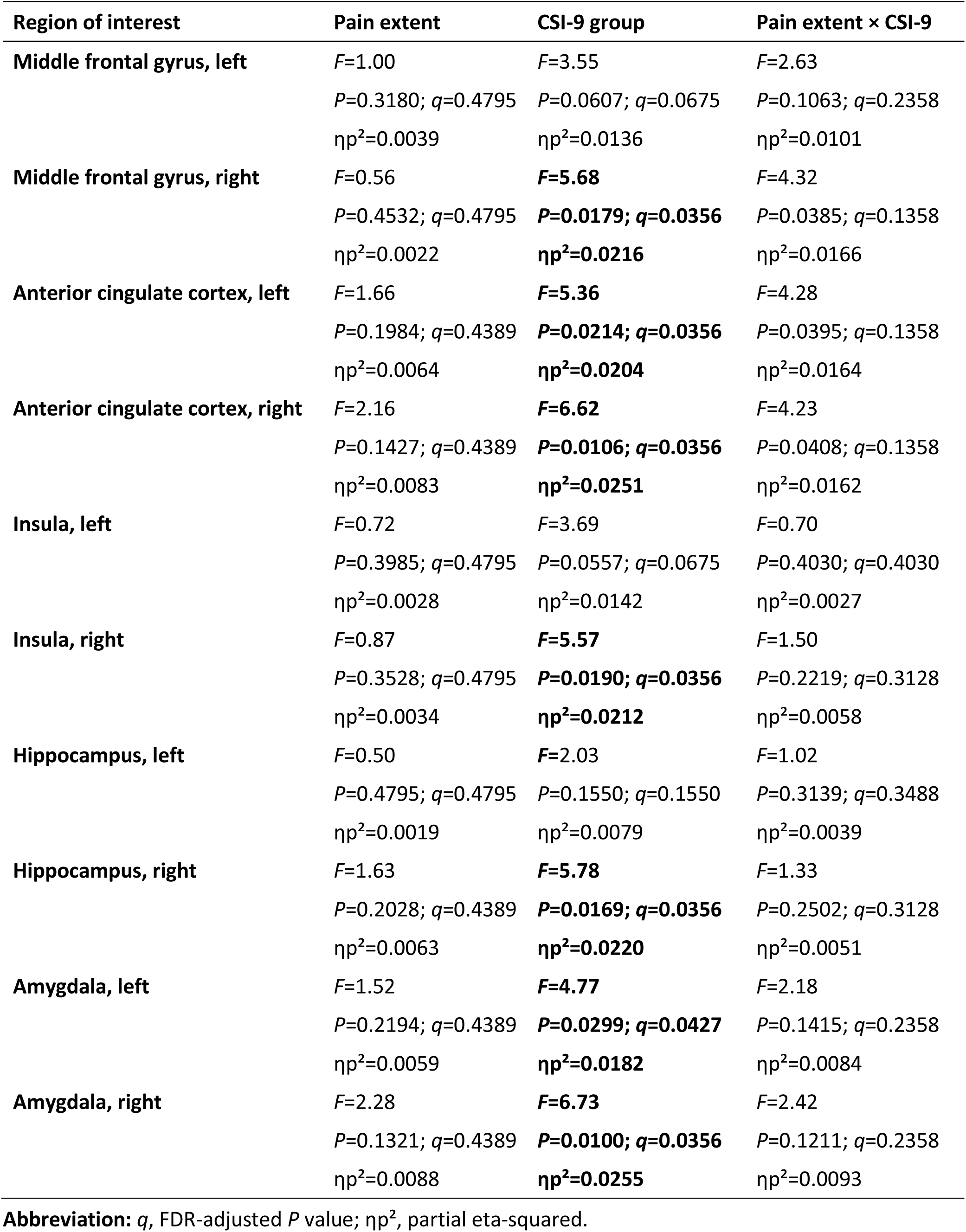
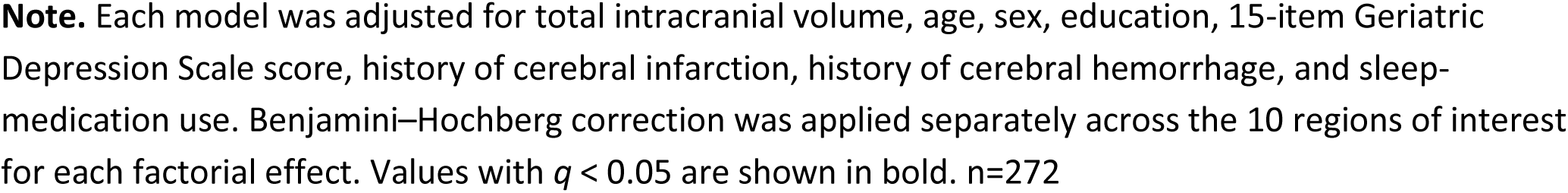
Regional gray-matter volume analyses according to pain extent and CSI-9 score.

| Region of interest | Pain extent | CSI-9 group | Pain extent × CSI-9 |
| --- | --- | --- | --- |
| **Middle frontal gyrus, left** | *F*=1.00<br>*P*=0.3180; *q*=0.4795<br>$\eta p^2$=0.0039 | *F*=3.55<br>*P*=0.0607; *q*=0.0675<br>$\eta p^2$=0.0136 | *F*=2.63<br>*P*=0.1063; *q*=0.2358<br>$\eta p^2$=0.0101 |
| **Middle frontal gyrus, right** | *F*=0.56<br>*P*=0.4532; *q*=0.4795<br>$\eta p^2$=0.0022 | ***F*=5.68**<br>***P*=0.0179; *q*=0.0356**<br>**$\eta p^2$=0.0216** | *F*=4.32<br>*P*=0.0385; *q*=0.1358<br>$\eta p^2$=0.0166 |
| **Anterior cingulate cortex, left** | *F*=1.66<br>*P*=0.1984; *q*=0.4389<br>$\eta p^2$=0.0064 | ***F*=5.36**<br>***P*=0.0214; *q*=0.0356**<br>**$\eta p^2$=0.0204** | *F*=4.28<br>*P*=0.0395; *q*=0.1358<br>$\eta p^2$=0.0164 |
| **Anterior cingulate cortex, right** | *F*=2.16<br>*P*=0.1427; *q*=0.4389<br>$\eta p^2$=0.0083 | ***F*=6.62**<br>***P*=0.0106; *q*=0.0356**<br>**$\eta p^2$=0.0251** | *F*=4.23<br>*P*=0.0408; *q*=0.1358<br>$\eta p^2$=0.0162 |
| **Insula, left** | *F*=0.72<br>*P*=0.3985; *q*=0.4795<br>$\eta p^2$=0.0028 | *F*=3.69<br>*P*=0.0557; *q*=0.0675<br>$\eta p^2$=0.0142 | *F*=0.70<br>*P*=0.4030; *q*=0.4030<br>$\eta p^2$=0.0027 |
| **Insula, right** | *F*=0.87<br>*P*=0.3528; *q*=0.4795<br>$\eta p^2$=0.0034 | ***F*=5.57**<br>***P*=0.0190; *q*=0.0356**<br>**$\eta p^2$=0.0212** | *F*=1.50<br>*P*=0.2219; *q*=0.3128<br>$\eta p^2$=0.0058 |
| **Hippocampus, left** | *F*=0.50<br>*P*=0.4795; *q*=0.4795<br>$\eta p^2$=0.0019 | *F*=2.03<br>*P*=0.1550; *q*=0.1550<br>$\eta p^2$=0.0079 | *F*=1.02<br>*P*=0.3139; *q*=0.3488<br>$\eta p^2$=0.0039 |
| **Hippocampus, right** | *F*=1.63<br>*P*=0.2028; *q*=0.4389<br>$\eta p^2$=0.0063 | ***F*=5.78**<br>***P*=0.0169; *q*=0.0356**<br>**$\eta p^2$=0.0220** | *F*=1.33<br>*P*=0.2502; *q*=0.3128<br>$\eta p^2$=0.0051 |
| **Amygdala, left** | *F*=1.52<br>*P*=0.2194; *q*=0.4389<br>$\eta p^2$=0.0059 | ***F*=4.77**<br>***P*=0.0299; *q*=0.0427**<br>**$\eta p^2$=0.0182** | *F*=2.18<br>*P*=0.1415; *q*=0.2358<br>$\eta p^2$=0.0084 |
| **Amygdala, right** | *F*=2.28<br>*P*=0.1321; *q*=0.4389<br>$\eta p^2$=0.0088 | ***F*=6.73**<br>***P*=0.0100; *q*=0.0356**<br>**$\eta p^2$=0.0255** | *F*=2.42<br>*P*=0.1211; *q*=0.2358<br>$\eta p^2$=0.0093 |

**Abbreviation:** *q*, FDR-adjusted *P* value; $\eta p^2$ , partial eta-squared.
**Note.** Each model was adjusted for total intracranial volume, age, sex, education, 15-item Geriatric Depression Scale score, history of cerebral infarction, history of cerebral hemorrhage, and sleep-medication use. Benjamini–Hochberg correction was applied separately across the 10 regions of interest for each factorial effect. Values with $q < 0.05$ are shown in bold. $n=272$

### 3.5 Sensitivity and Robustness Analyses

#### 3.5.1 TMT-B analyses

The pain extent × CSI-9 interaction remained statistically significant when a CSI-9 threshold of ≥16 was used (*P*=0.0323), after additional adjustment for cerebral small-vessel-disease markers (*P*=0.0072), with log-transformed TMT-B completion time (*P*=0.0476), and when cognitive performance was expressed as TMT-B minus TMT-A (p=0.0071) or as the TMT-B/TMT-A ratio (*P*=0.0135).

The interaction did not reach statistical significance when CSI-9 score was modeled continuously (*P*=0.0568), although this estimate was close to the conventional significance threshold and was directionally compatible with the primary categorical analysis. In contrast, the interaction was more clearly attenuated when both painful-region count and CSI-9 score were modeled continuously (*P*=0.8910), when multisite pain was defined as ≥3 regions (*P*=0.1616), or in the generalized estimating-equation model with empirical standard errors (*P*=0.1107). These results indicate that the interaction was supported by several, but not all, alternative model specifications (Table S3).

#### 3.5.2 Regional gray-matter volume analyses

When CSI-9 score was modeled continuously, no regional association remained significant after FDR correction; the smallest q value was observed for the right insula (q=0.0726). Using the alternative CSI-9 cutoff of ≥16, the CSI-9 main effect remained significant after FDR correction in the bilateral middle frontal gyri, bilateral anterior cingulate cortices, bilateral insulae, right hippocampus, and right amygdala. No interaction was significant after FDR correction in this analysis.

When multisite pain was defined as ≥3 regions, no pain-extent, CSI-9, or interaction effect remained significant after FDR correction. After additional adjustment for cerebral small-vessel-disease markers, the CSI-9 main effect remained significant after FDR correction in the right middle frontal gyrus, bilateral anterior cingulate cortices, right insula, right hippocampus, and right amygdala, whereas no interaction survived correction (Tables S5A–S5D).

#### 3.5.3 Contextual Comparison With Participants Reporting No Current Pain

The contextual TMT-B analysis included 298 participants reporting no current pain and the 272 participants in the four chronic-pain groups. The overall five-group effect was significant (*F*[4,555]=5.07, *P*=0.0005).

The adjusted mean TMT-B completion times were 124.26 seconds (95% CI, 117.60–130.92) in participants with no current pain, 121.19 seconds (95% CI, 106.61–135.78) in the single-site/lower CSI-9 group, 174.55 seconds (95% CI, 149.60–199.51) in the single-site/higher CSI-9 group, 116.40 seconds (95% CI, 106.58–126.22) in the multisite/lower CSI-9 group, and 121.76 seconds (95% CI, 111.37–132.15) in the multisite/higher CSI-9 group. Only the single-site/higher CSI-9 group had a significantly longer completion time than the no-current-pain group after Dunnett adjustment (p=0.0004); none of the other chronic-pain groups differed significantly from the no-current-pain group (Table S6).

#### 3.5.4 Leave-One-Out and Ceiling-Effect Analyses

In the leave-one-out analysis of the 14 participants in the single-site/higher CSI-9 group, the pain extent × CSI-9 interaction remained statistically significant in 13 of the 14 analyses. Interaction p values ranged from 0.0027 to 0.1492. When the participant with a TMT-B completion time of 445 seconds was omitted, the interaction was attenuated and was no longer statistically significant (*F*=2.09, *P*=0.1492), indicating that the primary result was influenced in part by this observation (Table S7).

The proportion of participants attaining the maximum MMSE score ranged from 14.0% to 24.5% across the four groups, whereas the corresponding proportion for the MoCA-J ranged from 4.5% to 7.1%. These distributions indicate a more prominent ceiling effect for the MMSE than for the MoCA-J (Table S8).

## 4. Discussion

In this community-based sample of older adults with chronic pain, pain extent and CSI-9 score showed different patterns of association with cognition and regional gray-matter volume. TMT-B completion time was longest in participants with single-site pain and a higher CSI-9 score, and the pain extent × CSI-9 interaction was statistically significant. No other cognitive outcome remained significant after correction for multiple testing. In contrast, regional gray-matter volumes were associated mainly with CSI-9 group: the higher CSI-9 group had smaller adjusted volumes in seven prespecified regions, whereas pain extent and the interaction were not significant after false-discovery-rate correction. Only the single-site/higher CSI-9 group showed slower TMT-B performance than participants reporting no current pain. These findings suggest that pain extent and CSI-9 score may represent partly distinct dimensions of chronic pain, while the small subgroup and mixed sensitivity results require caution.

### 4.1 Pain Extent and CSI-9 Score as Partly Distinct Dimensions

Pain extent describes the spatial distribution of pain, whereas the CSI-9 captures symptoms associated with central sensitization-related conditions. These dimensions may overlap, but they are not equivalent. Nociceptive, neuropathic, and nociplastic mechanisms can coexist within an individual, and pain distribution alone does not identify the mechanism sustaining pain (Fitzcharles et al., 2021; Kaplan et al., 2024). Consistent with this distinction, a higher CSI-9 score occurred in both single-site and multisite pain, and its association with TMT-B differed according to pain extent.

Slower TMT-B performance in the single-site/higher CSI-9 group should not be interpreted as evidence that single-site pain is inherently more centrally mediated than multisite pain. The group may have differed in underlying diagnoses, pain duration or intensity, medication use, or other unmeasured characteristics. Conversely, multisite pain may encompass heterogeneous combinations of musculoskeletal, neuropathic, visceral, and headache-related conditions. The contextual comparison indicated that the TMT-B finding was not solely a relative contrast with the multisite groups, because this group also performed more slowly than participants with no current pain. However, the interaction was attenuated in several alternative models and after omission of the participant with the longest TMT-B time. The observed pattern should therefore be considered hypothesis-generating rather than evidence of a stable categorical phenotype.

The CSI-9 is a brief patient-reported measure with established psychometric properties, but it does not directly quantify central nociceptive responsiveness (Nishigami et al., 2018). Associations between CSI scores and quantitative sensory testing vary across modalities and studies (Neblett et al., 2024). Accordingly, the present findings concern symptoms assessed by the CSI-9 and do not establish that central sensitization caused the cognitive or structural differences.

### 4.2 Cognitive Findings

The cognitive association was most evident for TMT-B, which depends on visual search, sequencing, sustained attention, processing speed, and set shifting. Chronic pain may compete with ongoing task demands, and meta-analytic evidence suggests a small attentional bias toward pain-related sensory information (Abudoush et al., 2023). Longitudinal and population-based studies have also associated chronic pain, particularly multisite pain, with cognitive decline and smaller hippocampal volume (Rouch et al., 2021; Zhao et al., 2023). These findings provide context for the present result, but attentional capture and competition for executive resources were not measured directly.

No secondary cognitive outcome remained significant after false-discovery-rate correction. Analyses using TMT-B minus TMT-A and the TMT-B/TMT-A ratio retained the interaction, suggesting that the finding was not explained entirely by simple visuomotor speed. Nevertheless, TMT-B reflects multiple cognitive and motor processes, the MMSE showed a greater ceiling effect than the MoCA-J, and the single-site/higher CSI-9 estimate was imprecise. The results therefore support slower TMT-B performance rather than a definitive or isolated executive impairment.

### 4.3 Regional Gray-Matter Volume Findings

Regional gray-matter volumes showed a main effect of CSI-9 group rather than pain extent or an interaction. The implicated regions—the middle frontal gyrus, anterior cingulate cortex, insula, hippocampus, and amygdala—participate in distributed systems supporting cognitive control, salience detection, affective processing, learning, and memory. The anterior cingulate cortex and insula are central to salience processing but are not specific to pain (Uddin, 2015).

The locations of the observed differences are broadly consistent with recent structural imaging literature. A preregistered meta-analysis found no common chronic-pain structural difference after stringent correction but identified subtle, spatially distributed alterations in exploratory analyses (Henn et al., 2023). A population-based study likewise reported lower gray-matter volume in the anterior cingulate cortex, insula, and hippocampus across several chronic pain conditions (Neumann et al., 2023). In the present study, effect sizes were small, and the results depended partly on how CSI-9 was modeled. Several associations remained significant using a cutoff of 16, whereas none survived false-discovery-rate correction when CSI-9 was treated continuously. This discrepancy may reflect nonlinearity, limited power, or categorization effects and does not establish a biologically discrete threshold.

The cognitive and structural patterns were not parallel. TMT-B showed an interaction between pain extent and CSI-9 group, whereas regional volumes were associated mainly with CSI-9 group. The findings therefore do not support a simple explanation in which smaller regional volume accounts for slower TMT-B performance in the single-site/higher CSI-9 group. Cognitive performance and brain structure may represent partly distinct correlates or may change on different timescales. Cerebral small-vessel disease is a relevant competing explanation in older adults because white-matter hyperintensities and lacunes are associated with cognitive decline (Jansma et al., 2024). Additional adjustment for measured small-vessel-disease markers did not materially change the TMT-B interaction and preserved several CSI-9-related regional associations. These findings reduce, but do not eliminate, the possibility that cerebral small-vessel disease confounded the observed associations of pain phenotype with cognitive performance or regional gray-matter volume.

### 4.4 Clinical and Conceptual Implications

The findings do not establish a new diagnostic entity. Rather, they indicate that pain extent and central sensitization-related symptoms should not be treated as interchangeable. Pain may remain spatially limited even when the CSI-9 score is high, while multisite pain may arise from several peripheral or mixed mechanisms without uniformly high CSI-9 scores. Assessing both dimensions may therefore describe chronic pain heterogeneity more informatively than either alone.

The CSI-9 may be useful as one component of a broader clinical assessment, but the present results do not support using a cutoff as a diagnostic test for central sensitization, a marker of brain atrophy, or a stand-alone basis for treatment selection. Clinical applications require replication, etiologic characterization of pain, and prospective evaluation of outcomes and treatment response.

### 4.5 Strengths and Limitations

Strengths include the community-based older sample and the factorial design separating pain extent from CSI-9 score. The analysis used a prespecified primary cognitive outcome, multiple-testing correction, covariate-adjusted marginal means, alternative exposure and outcome definitions, additional small-vessel-disease adjustment, comparison with participants reporting no current pain, and leave-one-out analysis of the smallest group.

The current study involved several limitations that should be noted. First, the cross-sectional design precludes conclusions about temporal order or causality; smaller volumes cannot be interpreted as pain-related atrophy. Second, the single-site/higher CSI-9 group included only 14 participants, and the TMT-B interaction was influenced by one participant with a particularly long completion time and was not significant in several sensitivity analyses. Third, the CSI-9 is self-reported, and quantitative sensory testing was unavailable. Fourth, pain extent was based on seven broad regions and does not correspond to chronic widespread pain criteria. Etiologic diagnoses were unavailable, limiting assessment of diagnostic heterogeneity.

Fifth, MRI was acquired at 0.3 T. Volumes from the same scanner have shown good agreement with 3-T measurements for major brain structures, supporting its use in a community setting (Murata et al., 2022). Nevertheless, lower signal-to-noise ratio and spatial resolution may have reduced sensitivity to subtle differences, and the region-of-interest approach could not detect effects elsewhere. Sixth, residual confounding by pain diagnosis, medication, physical activity, cardiometabolic disease, anxiety, sleep quality, socioeconomic factors, or other age-related pathology remains possible. Finally, participants able to attend repeated on-site assessments and MRI may have been healthier than the broader older population.

In conclusion, pain extent and CSI-9 score appeared to represent partly distinct dimensions among older adults with chronic pain. A higher CSI-9 score was associated with slower TMT-B performance primarily in participants with single-site pain, whereas smaller regional gray-matter volumes were associated mainly with CSI-9 group. The differing cognitive and structural patterns, small subgroup size, and attenuation in several sensitivity analyses warrant cautious interpretation. Larger longitudinal studies incorporating pain diagnoses, quantitative sensory testing, and repeated cognitive and neuroimaging assessments are needed.

## Acknowledgments

This work was supported by the Health Labour Sciences Research Grant (Grant Number 25FG1001); JSPS KAKENHI Grants (Grant Numbers JP18H03184 and JP24K21147); the MEXT-Supported Program for the Strategic Research Foundation at Private Universities, 2014-2018 (Grant Number S1411006); Grant-in-Aid for Special Research in Subsidies for ordinary expenses of private schools from the Promotion and Mutual Aid Corporation for Private Schools of Japan; Mizuno Sports Promotion Foundation; and the Mitsui Life Social Welfare Foundation. We acknowledge the use of ChatGPT (GPT-5.5 model; OpenAI, San Francisco, California), Perplexity (Perplexity AI, Inc., San Francisco, California), using the Gemini 3 Pro model (Google, Mountain View, California), and NotebookLM (Google, Mountain View, California) on January 2, 2026. These tools assisted with English editing and improving readability (e.g., refining phrasing and sentence structure) in portions of the introduction, methods, and discussion, and with preparing the graphical abstract. All AI-assisted text was reviewed and edited by the authors, who take full responsibility for the integrity and accuracy of the content.

## Ethical Statement

The study protocol adhered to the Declaration of Helsinki of 1975, as revised in 2024, and received approval from the Research Ethics Committee, Faculty of Medicine, Juntendo University in September 2015 (first approval no. 2015061, and the latest revised version no. M15-0057-M13). Written informed consent was obtained from every participant prior to enrollment, and they were notified that they had the right to withdraw from the study at any time.

## Consent statement

All participants provided written informed consent.

## Data availability statement

The datasets generated during and/or analyzed during the current study are not publicly available due to privacy restrictions protecting participant identity but are available from the corresponding author on reasonable request, subject to ethical approval. Researchers interested in accessing the data should contact the data manager, Dr. Yoshifumi Tamura, at. Requests will be considered on an individual basis, taking into account ethical considerations and study requirements.

## Author Contribution Statement

K.Y.: Conceptualization; Methodology; Investigation; Formal analysis; Writing - original draft; Writing - review & editing; Funding acquisition. H.T.: Investigation; Writing - review & editing; Funding acquisition. H.N.: Investigation; Writing - review & editing. H.K.: Investigation; Writing - review & editing. Y.T.: Investigation; Writing - review & editing; Funding acquisition. K.T.: Writing - review & editing. K.K.: Writing - review & editing.

---

**Table S1.** Distribution of painful regions across the four chronic-pain phenotype groups.

| **Pain region** | **Single-site**<br>**/ Lower CSI-9**<br>**(n=43)** | **Single-site**<br>**/ Higher CSI-9**<br>**(n=14)** | **Multisite**<br>**/ Lower CSI-9**<br>**(n=110)** | **Multisite**<br>**/ Higher CSI-9**<br>**(n=105)** |
| --- | --- | --- | --- | --- |
| **Left upper extremity** | 3 (7.0) | 3 (21.4) | 41 (37.3) | 47 (44.8) |
| **Right upper extremity** | 8 (18.6) | 1 (7.1) | 48 (43.6) | 62 (59.0) |
| **Left lower extremity** | 6 (13.9) | 1 (7.1) | 65 (59.1) | 70 (66.7) |
| **Right lower extremity** | 12 (27.9) | 2 (14.3) | 70 (63.6) | 74 (70.5) |
| **Trunk** | 13 (30.2) | 5 (35.7) | 68 (61.8) | 85 (81.0) |
| **Head/craniofacial/throat** | 1 (2.3) | 2 (14.3) | 21 (19.1) | 35 (33.3) |
| **Visceral/pelvic region** | 0 (0) | 0 (0) | 10 (9.1) | 13 (12.4) |

**Abbreviation:** CSI-9, 9-item Central Sensitization Inventory.
**Note.** Values are n (%). Higher CSI-9 was defined as $\geq 12$ . Single-site pain was defined as exactly one of seven mapped pain regions; multisite pain was defined as $\geq 2$ regions. n=272

**Table S2.** Secondary cognitive outcomes: factorial analysis of covariance with False Discovery Rate correction.

| Outcome | Pain extent | CSI-9 group | Pain extent × CSI-9 |
| --- | --- | --- | --- |
| TMT-A, sec | F=0.90<br>p=0.3440; q=0.5160<br>$\eta^2$=0.0035 | F=0.00<br>p=0.9814; q=0.9814<br>$\eta^2$<0.0001 | F=0.63<br>p=0.4275; q=0.5609<br>$\eta^2$=0.0024 |
| MMSE total score | F=0.96<br>p=0.3288; q=0.5160<br>$\eta^2$=0.0037 | F=0.12<br>p=0.7256; q=0.9814<br>$\eta^2$=0.0005 | F=2.33<br>p=0.1284; q=0.3706<br>$\eta^2$=0.0089 |
| MoCA-J total score | F=2.07<br>p=0.1510; q=0.5160<br>$\eta^2$=0.0080 | F=0.04<br>p=0.8457; q=0.9814<br>$\eta^2$=0.0001 | F=0.34<br>p=0.5609; q=0.5609<br>$\eta^2$=0.0013 |
| MoCA-J attention | F=1.70<br>p=0.1935; q=0.5160<br>$\eta^2$=0.0065 | F=0.30<br>p=0.5826; q=0.9814<br>$\eta^2$=0.0012 | F=0.34<br>p=0.5590; q=0.5609<br>$\eta^2$=0.0013 |
| MoCA-J visuospatial/executive | F=0.43<br>p=0.5124; q=0.5227<br>$\eta^2$=0.0017 | F=0.71<br>p=0.3989; q=0.9814<br>$\eta^2$=0.0028 | F=6.66<br>p=0.0104; q=0.0625<br>$\eta^2$=0.0252 |
| MoCA-J abstraction | F=0.41<br>p=0.5227; q=0.5227<br>$\eta^2$=0.0016 | F=0.43<br>p=0.5115; q=0.9814<br>$\eta^2$=0.0017 | F=1.76<br>p=0.1853; q=0.3706<br>$\eta^2$=0.0068 |

**Abbreviation:** CSI-9, 9-item Central Sensitization Inventory; TMT-A, Trail Making Test Part A; MMSE, Mini-Mental State Examination; MoCA-J, Japanese version of the Montreal Cognitive Assessment; q, FDR-adjusted p value; $\eta^2$ , partial eta-squared.
**Note.** Models were adjusted for age, sex, education, 15-item Geriatric Depression Scale score, history of cerebral infarction, history of cerebral hemorrhage, and sleep-medication use. Benjamini–Hochberg correction was applied separately across the six outcomes for each effect.

**Table S3.** Sensitivity and robustness analyses for Trail Making Test B.

| Analysis | Pain-extent term | CSI-9 term | Interaction term |
| --- | --- | --- | --- |
| Continuous CSI-9 | **F=8.27; p=0.0044** | **F=7.35; p=0.0072** | F=3.66; p=0.0568 |
| Continuous painful-region count × continuous CSI-9 | F=2.55; p=0.1112 | **F=3.98; p=0.0471** | F=0.02; p=0.8910 |
| CSI-9 threshold ≥16 | **F=8.25; p=0.0044** | **F=7.95; p=0.0052** | **F=4.63; p=0.0323** |
| Multisite threshold ≥3 regions | F=1.77; p=0.1842 | **F=4.01; p=0.0463** | F=1.97; p=0.1616 |
| Additional cerebral small-vessel-disease adjustment | **F=10.30; p=0.0015** | **F=8.88; p=0.0032** | **F=7.35; p=0.0072** |
| Log-transformed TMT-B | **F=5.61; p=0.0186** | **F=5.54; p=0.0193** | **F=3.96; p=0.0476** |
| TMT-B minus TMT-A | **F=14.19; p=0.0002** | **F=11.85; p=0.0007** | **F=7.37; p=0.0071** |
| TMT-B/TMT-A ratio | **F=14.57; p=0.0002** | **F=10.87; p=0.0011** | **F=6.19; p=0.0135** |
| GEE with empirical standard errors | **Wald $\chi^2=3.44$; p=0.0636** | **Wald $\chi^2=4.68$; p=0.0305** | Wald $\chi^2=2.54$; p=0.1107 |
| Leave-one-out analysis of the single-site/higher CSI-9 group | — | — | **13/14 analyses p<0.05; p range 0.0027–0.1492** |

**Abbreviation:** CSI-9, 9-item Central Sensitization Inventory; TMT-A, Trail Making Test part A; TMT-B, Trail Making Test part B; GEE, Generalized Estimating Equations.
**Note.** F statistics are shown except for the GEE analysis, which reports Wald $\chi^2$ statistics. For the continuous painful-region-count model, the first term is the continuous number of painful regions. All models used the same covariates as the primary analysis; the small-vessel-disease model additionally included periventricular hyperintensity, deep/subcortical white-matter hyperintensity, and lacune count. Values with $p < 0.05$ are shown in bold.

**Table S4.** Covariate-adjusted regional gray-matter volumes across the four phenotype groups.

| Region of interest | Single-site<br>/ Lower CSI-9<br>(n=43) | Single-site<br>/ Higher CSI-9<br>(n=14) | Multisite<br>/ Lower CSI-9<br>(n=110) | Multisite<br>/ Higher CSI-9<br>(n=105) |
| --- | --- | --- | --- | --- |
| Middle frontal gyrus, left | 10.311 | 9.499 | 10.172 | 10.090 |
| Middle frontal gyrus, right | 11.452 | 10.414 | 11.152 | 11.055 |
| Anterior cingulate cortex, left | 4.332 | 3.905 | 4.258 | 4.224 |
| Anterior cingulate cortex, right | 3.829 | 3.441 | 3.782 | 3.729 |
| Insula, left | 5.315 | 5.023 | 5.316 | 5.195 |
| Insula, right | 5.014 | 4.642 | 4.985 | 4.859 |
| Hippocampus, left | 2.726 | 2.603 | 2.711 | 2.687 |
| Hippocampus, right | 2.711 | 2.526 | 2.718 | 2.648 |
| Amygdala, left | 0.813 | 0.756 | 0.809 | 0.797 |
| Amygdala, right | 0.902 | 0.834 | 0.901 | 0.883 |

**Abbreviation:** CSI-9, 9-item Central Sensitization Inventory.
**Note.** Values are estimated marginal means in cm<sup>3</sup>. Models were adjusted for total intracranial volume, age, sex, education, 15-item Geriatric Depression Scale score, history of cerebral infarction, history of cerebral hemorrhage, and sleep-medication use.

**Table S5A.** Regional gray-matter volumes with CSI-9 modeled continuously.

| Region of interest | Pain extent | Continuous CSI-9 | Pain extent × CSI-9 |
| --- | --- | --- | --- |
| **Middle frontal gyrus, left** | F=0.46<br>p=0.4987; q=0.5975 | F=2.52<br>p=0.1139; q=0.1183 | F=0.34<br>p=0.5606; q=0.7965 |
| **Middle frontal gyrus, right** | F=0.10<br>p=0.7506; q=0.7506 | F=3.82<br>p=0.0518; q=0.1029 | F=0.59<br>p=0.4449; q=0.7965 |
| **Anterior cingulate cortex, left** | F=0.90<br>p=0.3429; q=0.5916 | F=3.87<br>p=0.0503; q=0.1029 | F=1.11<br>p=0.2920; q=0.7965 |
| **Anterior cingulate cortex, right** | F=1.11<br>p=0.2927; q=0.5916 | F=3.19<br>p=0.0754; q=0.1077 | F=1.17<br>p=0.2802; q=0.7965 |
| **Insula, left** | F=0.67<br>p=0.4141; q=0.5916 | F=4.93<br>p=0.0273; q=0.1029 | F=0.02<br>p=0.8830; q=0.8830 |
| **Insula, right** | F=0.76<br>p=0.3855; q=0.5916 | F=7.32<br>p=0.0073; q=0.0726 | F=0.14<br>p=0.7120; q=0.7965 |
| **Hippocampus, left** | F=0.38<br>p=0.5378; q=0.5975 | F=2.64<br>p=0.1052; q=0.1183 | F=0.13<br>p=0.7169; q=0.7965 |
| **Hippocampus, right** | F=1.20<br>p=0.2751; q=0.5916 | F=3.66<br>p=0.0569; q=0.1029 | F=0.34<br>p=0.5621; q=0.7965 |
| **Amygdala, left** | F=0.99<br>p=0.3212; q=0.5916 | F=2.46<br>p=0.1183; q=0.1183 | F=0.97<br>p=0.3246; q=0.7965 |
| **Amygdala, right** | F=1.71<br>p=0.1927; q=0.5916 | F=3.52<br>p=0.0617; q=0.1029 | F=1.49<br>p=0.2237; q=0.7965 |

**Abbreviation:** CSI-9, 9-item Central Sensitization Inventory; q, FDR-adjusted p value.
**Note.** Benjamini–Hochberg correction was applied separately across the 10 regions of interest for each effect.

**Table S5B.** Regional gray-matter volumes using CSI-9 ≥16.

| Region of interest | Pain extent | CSI-9 $\geq 16$ | Interaction |
| --- | --- | --- | --- |
| Middle frontal gyrus, left | F=1.08<br>p=0.2991; q=0.3323 | F=5.49<br>**p=0.0198; q=0.0248** | F=1.96<br>p=0.1630; q=0.1962 |
| Middle frontal gyrus, right | F=1.15<br>p=0.2846; q=0.3323 | F=8.88<br>**p=0.0032; q=0.0100** | F=4.25<br>p=0.0403; q=0.1009 |
| Anterior cingulate cortex, left | F=3.05<br>p=0.0818; q=0.2282 | F=8.74<br>**p=0.0034; q=0.0100** | F=5.62<br>p=0.0185; q=0.0616 |
| Anterior cingulate cortex, right | F=3.72<br>p=0.0548; q=0.2282 | F=8.44<br>**p=0.0040; q=0.0100** | F=6.08<br>p=0.0143; q=0.0616 |
| Insula, left | F=1.15<br>p=0.2839; q=0.3323 | F=6.35<br>**p=0.0124; q=0.0206** | F=1.81<br>p=0.1798; q=0.1962 |
| Insula, right | F=1.57<br>p=0.2120; q=0.3323 | F=10.83<br>**p=0.0011; q=0.0100** | F=3.12<br>p=0.0785; q=0.1148 |
| Hippocampus, left | F=0.93<br>p=0.3363; q=0.3363 | F=3.25<br>p=0.0724; q=0.0804 | F=1.68<br>p=0.1962; q=0.1962 |
| Hippocampus, right | F=2.87<br>p=0.0913; q=0.2282 | F=7.01<br>**p=0.0086; q=0.0172** | F=3.85<br>p=0.0508; q=0.1016 |
| Amygdala, left | F=2.19<br>p=0.1399; q=0.2797 | F=2.29<br>p=0.1314; q=0.1314 | F=3.08<br>p=0.0804; q=0.1148 |
| Amygdala, right | F=4.35<br>p=0.0379; q=0.2282 | F=5.55<br>**p=0.0192; q=0.0248** | F=5.98<br>p=0.0151; q=0.0616 |

**Abbreviation:** CSI-9, 9-item Central Sensitization Inventory; q, FDR-adjusted p value.
**Note.** Benjamini–Hochberg correction was applied separately across the 10 regions of interest for each effect. Values with $q < 0.05$ are shown in bold.

**Table S5C.** Regional gray-matter volumes defining multisite pain as ≥3 regions.

| Region of interest | Pain extent<br>($\geq 3$ regions) | CSI-9 group | Interaction |
| --- | --- | --- | --- |
| Middle frontal gyrus<br>, left | F=0.75<br>p=0.3878; q=0.3878 | F=1.48<br>p=0.2246; q=0.2246 | F<0.01<br>p=0.9923; q=0.9923 |
| Middle frontal gyrus<br>, right | F=1.47<br>p=0.2261; q=0.3634 | F=3.18<br>p=0.0756; q=0.1080 | F=0.81<br>p=0.3679; q=0.9198 |
| Anterior cingulate cortex<br>, left | F=1.31<br>p=0.2544; q=0.3634 | F=2.78<br>p=0.0969; q=0.1211 | F=1.58<br>p=0.2094; q=0.9198 |
| Anterior cingulate cortex<br>, right | F=1.64<br>p=0.2012; q=0.3634 | F=3.79<br>p=0.0527; q=0.1054 | F=0.86<br>p=0.3551; q=0.9198 |
| Insula, left | F=1.61<br>p=0.2051; q=0.3634 | F=4.19<br>p=0.0416; q=0.1040 | F=0.17<br>p=0.6816; q=0.9737 |
| Insula, right | F=1.75<br>p=0.1868; q=0.3634 | F=5.42<br>p=0.0207; q=0.1035 | F=0.28<br>p=0.5984; q=0.9737 |
| Hippocampus, left | F=1.71<br>p=0.1925; q=0.3634 | F=2.06<br>p=0.1526; q=0.1695 | F=0.98<br>p=0.3243; q=0.9198 |
| Hippocampus, right | F=2.39<br>p=0.1233; q=0.3634 | F=5.67<br>p=0.0180; q=0.1035 | F=0.03<br>p=0.8683; q=0.9923 |
| Amygdala, left | F=1.08<br>p=0.3002; q=0.3753 | F=3.34<br>p=0.0687; q=0.1080 | F=0.50<br>p=0.4805; q=0.9609 |
| Amygdala, right | F=0.77<br>p=0.3807; q=0.3878 | F=4.33<br>p=0.0383; q=0.1040 | F=0.00<br>p=0.9448; q=0.9923 |

**Abbreviation:** CSI-9, 9-item Central Sensitization Inventory; q, FDR-adjusted p value.
**Note.** Benjamini–Hochberg correction was applied separately across the 10 regions of interest for each effect.

**Table S5D.** Regional gray-matter volumes additionally adjusted for cerebral small-vessel-disease markers.

| Region of interest | Pain extent | CSI-9 group | Interaction |
| --- | --- | --- | --- |
| Middle frontal gyrus , left | F=1.02<br>p=0.3128; q=0.4808 | F=3.72<br>p=0.0549; q=0.0686 | F=2.61<br>p=0.1072; q=0.2357 |
| Middle frontal gyrus , right | F=0.55<br>p=0.4610; q=0.5116 | **F=5.80**<br>**p=0.0168; q=0.0451** | F=4.32<br>p=0.0387; q=0.1409 |
| Anterior cingulate cortex , left | F=1.58<br>p=0.2106; q=0.4797 | **F=5.06**<br>**p=0.0254; q=0.0451** | F=4.23<br>p=0.0406; q=0.1409 |
| Anterior cingulate cortex , right | F=1.98<br>p=0.1610; q=0.4797 | **F=6.78**<br>**p=0.0098; q=0.0451** | F=4.17<br>p=0.0423; q=0.1409 |
| Insula, left | F=0.76<br>p=0.3846; q=0.4808 | F=2.91<br>p=0.0893; q=0.0993 | F=0.71<br>p=0.4012; q=0.4012 |
| Insula, right | F=0.92<br>p=0.3389; q=0.4808 | **F=5.00**<br>**p=0.0262; q=0.0451** | F=1.52<br>p=0.2193; q=0.3094 |
| Hippocampus, left | F=0.43<br>p=0.5116; q=0.5116 | F=1.66<br>p=0.1994; q=0.1994 | F=1.01<br>p=0.3168; q=0.3520 |
| Hippocampus, right | F=1.68<br>p=0.1967; q=0.4797 | **F=4.94**<br>**p=0.0271; q=0.0451** | F=1.34<br>p=0.2475; q=0.3094 |
| Amygdala, left | F=1.39<br>p=0.2398; q=0.4797 | F=4.01<br>p=0.0462; q=0.0661 | F=2.18<br>p=0.1414; q=0.2357 |
| Amygdala, right | F=2.15<br>p=0.1440; q=0.4797 | **F=6.11**<br>**p=0.0141; q=0.0451** | F=2.41<br>p=0.1219; q=0.2357 |

**Abbreviation:** CSI-9, 9-item Central Sensitization Inventory; q, FDR-adjusted p value.
**Note.** Benjamini–Hochberg correction was applied separately across the 10 regions of interest for each effect. Values with $q < 0.05$ are shown in bold.

**Table S6.**
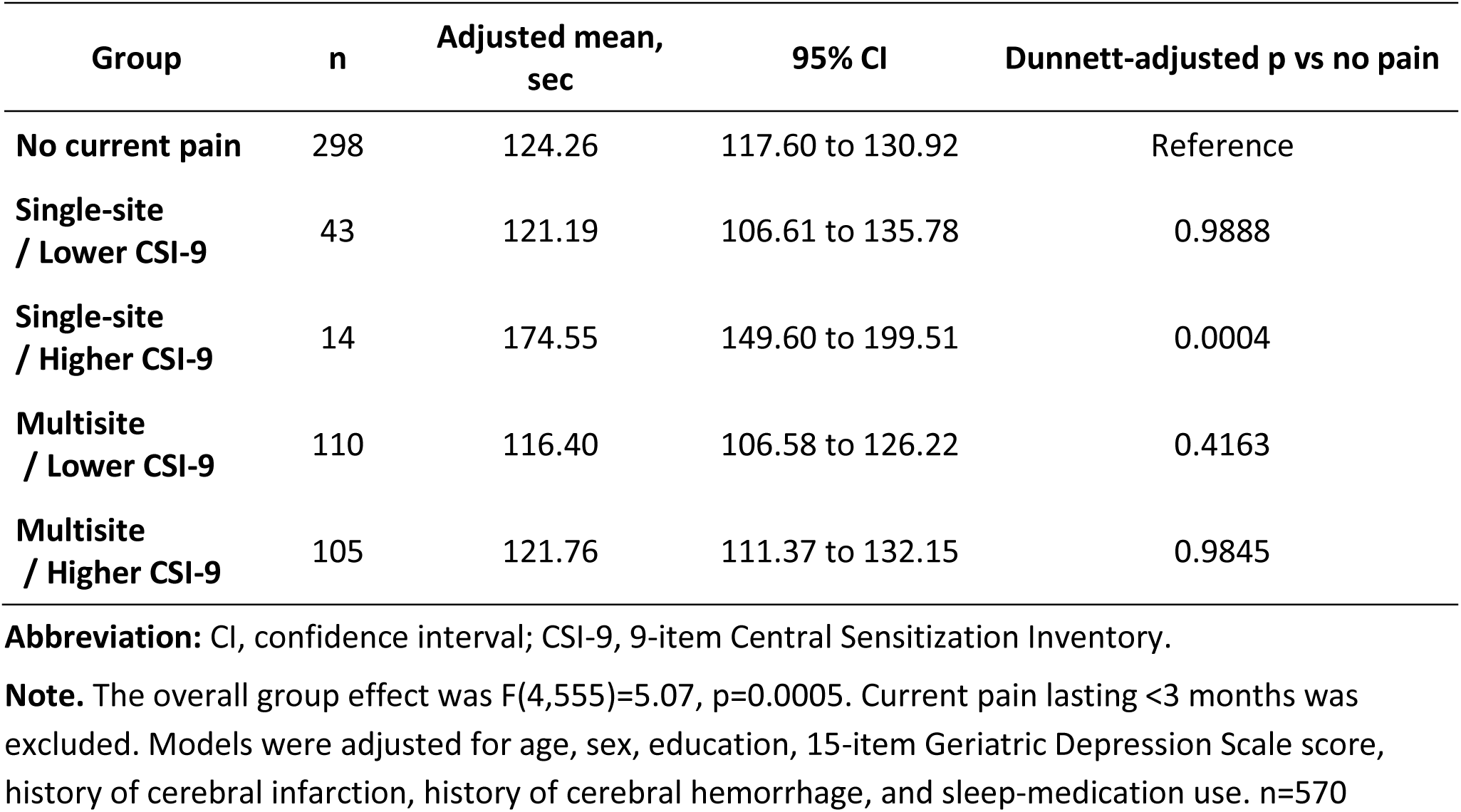
Contextual comparison of Trail Making Test Part B performance with participants reporting no current pain.

| Group | n | Adjusted mean, sec | 95% CI | Dunnett-adjusted p vs no pain |
| --- | --- | --- | --- | --- |
| No current pain | 298 | 124.26 | 117.60 to 130.92 | Reference |
| Single-site / Lower CSI-9 | 43 | 121.19 | 106.61 to 135.78 | 0.9888 |
| Single-site / Higher CSI-9 | 14 | 174.55 | 149.60 to 199.51 | 0.0004 |
| Multisite / Lower CSI-9 | 110 | 116.40 | 106.58 to 126.22 | 0.4163 |
| Multisite / Higher CSI-9 | 105 | 121.76 | 111.37 to 132.15 | 0.9845 |

**Abbreviation:** CI, confidence interval; CSI-9, 9-item Central Sensitization Inventory.
**Note.** The overall group effect was $F(4,555)=5.07$ , $p=0.0005$ . Current pain lasting <3 months was excluded. Models were adjusted for age, sex, education, 15-item Geriatric Depression Scale score, history of cerebral infarction, history of cerebral hemorrhage, and sleep-medication use. n=570

**Table S7.** Leave-one-out analysis of the pain extent × CSI-9 interaction for Trail Making Test Part B.

| Omission order | Omitted TMT-B value, sec | F for interaction | p |
| --- | --- | --- | --- |
| 1 | 80 | 9.07 | 0.0029 |
| 2 | 144 | 7.83 | 0.0055 |
| 3 | 325 | 5.05 | 0.0254 |
| 4 | 198 | 6.50 | 0.0113 |
| 5 | 220 | 6.63 | 0.0106 |
| 6 | 277 | 5.95 | 0.0154 |
| 7 | 93 | 8.93 | 0.0031 |
| 8 | 192 | 6.27 | 0.0129 |
| 9 | 115 | 8.82 | 0.0033 |
| 10 | 445 | 2.09 | 0.1492 |
| 11 | 167 | 7.45 | 0.0068 |
| 12 | 91 | 9.04 | 0.0029 |
| 13 | 99 | 8.81 | 0.0033 |
| 14 | 119 | 9.18 | 0.0027 |

**Abbreviation:** CSI-9, 9-item Central Sensitization Inventory.
**Note.** Participant identifiers are intentionally omitted. The interaction remained significant in 13 of 14 analyses; the result was attenuated when the participant with Trail Making Test Part B =445 sec was omitted (p=0.1492).

**Table S8.** Ceiling-effect checks for MMSE and MoCA-J.

| Group | n | MMSE<br>maximum, n | MMSE<br>maximum, % | MoCA-J<br>maximum, n | MoCA-J<br>maximum, % |
| --- | --- | --- | --- | --- | --- |
| Single-site /<br>Lower CSI-9 | 43 | 6 | 14.0 | 2 | 4.7 |
| Single-site /<br>Higher CSI-9 | 14 | 3 | 21.4 | 1 | 7.1 |
| Multisite /<br>Lower CSI-9 | 110 | 27 | 24.5 | 5 | 4.5 |
| Multisite /<br>Higher CSI-9 | 105 | 19 | 18.1 | 6 | 5.7 |

**Abbreviation:** CSI-9, 9-item Central Sensitization Inventory; MMSE, Mini-Mental State Examination; MoCA-J, Japanese version of the Montreal Cognitive Assessment.
**Note.** The maximum score was 30 for both MMSE and MoCA-J.

## References

Abudoush, A. N., Noureen, A., Panagioti, M., Poliakoff, E., Van Ryckeghem, D. M. L., Hodkinson, A., & Husain, N. (2023). What can we learn about selective attention processes in individuals with chronic pain using reaction time tasks? A systematic review and meta-analysis. Pain, 164(8), 1677–1692.

Ashburner, John. (2007). A fast diffeomorphic image registration algorithm. NeuroImage, 38(1), 95– 113.

Ashburner, John, & Friston, K. J. (2000). Voxel-based morphometry--the methods. NeuroImage, 11(6 Pt 1), 805–821.

Benjamini, Y., & Hochberg, Y. (1995). Controlling the false discovery rate: A practical and powerful approach to multiple testing. Journal of the Royal Statistical Society. Series B, Statistical Methodology, 57(1), 289–300.

Cohen, S. P., Vase, L., & Hooten, W. M. (2021). Chronic pain: an update on burden, best practices, and new advances. Lancet, 397(10289), 2082–2097.

Domenichiello, A. F., & Ramsden, C. E. (2019). The silent epidemic of chronic pain in older adults. Progress in Neuro-Psychopharmacology & Biological Psychiatry, 93, 284–290.

Fazekas, F., Chawluk, J. B., Alavi, A., Hurtig, H. I., & Zimmerman, R. A. (1987). MR signal abnormalities at 1.5 T in Alzheimer’s dementia and normal aging. AJR. American Journal of Roentgenology, 149(2), 351–356.

Fitzcharles, M.-A., Cohen, S. P., Clauw, D. J., Littlejohn, G., Usui, C., & Häuser, W. (2021). Nociplastic pain: towards an understanding of prevalent pain conditions. Lancet, 397(10289), 2098– 2110.

Fujiwara, Y., Suzuki, H., Yasunaga, M., Sugiyama, M., Ijuin, M., Sakuma, N., Inagaki, H., Iwasa, H., Ura, C., Yatomi, N., Ishii, K., Tokumaru, A. M., Homma, A., Nasreddine, Z., & Shinkai, S. (2010). Brief screening tool for mild cognitive impairment in older Japanese: validation of the Japanese version of the Montreal Cognitive Assessment. Geriatrics & Gerontology International, 10(3), 225–232.

Goldstein, F. C., Ashley, A. V., Miller, E., Alexeeva, O., Zanders, L., & King, V. (2014). Validity of the montreal cognitive assessment as a screen for mild cognitive impairment and dementia in African Americans. Journal of Geriatric Psychiatry and Neurology, 27(3), 199–203.

Henn, A. T., Larsen, B., Frahm, L., Xu, A., Adebimpe, A., Scott, J. C., Linguiti, S., Sharma, V., Basbaum, A. I., Corder, G., Dworkin, R. H., Edwards, R. R., Woolf, C. J., Habel, U., Eickhoff, S. B., Eickhoff, C. R., Wagels, L., & Satterthwaite, T. D. (2023). Structural imaging studies of patients with chronic pain: an anatomical likelihood estimate meta-analysis. Pain, 164(1), e10–e24.

Hoshino, H., Sasaki, N., Ide, K., Yamato, Y., Watanabe, Y., & Matsuyama, Y. (2022). Effect of central sensitization inventory on the number of painful sites and pain severity in a Japanese regional population cohort. Journal of Orthopaedic Science: Official Journal of the Japanese Orthopaedic Association, 27(4), 929–934.

Ideno, Y., Takayama, M., Hayashi, K., Takagi, H., & Sugai, Y. (2012). Evaluation of a Japanese version of the Mini-Mental State Examination in elderly persons. Geriatrics & Gerontology International, 12(2), 310–316.

Inoue, S., Hashizume, H., Murata, S., Oka, H., Kozaki, T., Minakata, K., Taiji, R., Teraguchi, M., Iwasaki, H., Tsutsui, S., Takami, M., Mure, K., Nakagawa, Y., Miyai, N., & Yamada, H. (2025). Association between central sensitization and multisite pain in the general population: A cross-sectional analysis of The Wakayama Health Promotion Study. Journal of Orthopaedic Science: Official Journal of the Japanese Orthopaedic Association, 30(6), 1186–1192.

Jansma, A., de Bresser, J., Schoones, J. W., van Heemst, D., & Akintola, A. A. (2024). Sporadic cerebral small vessel disease and cognitive decline in healthy older adults: A systematic review and meta-analysis. Journal of Cerebral Blood Flow and Metabolism: Official Journal of the International Society of Cerebral Blood Flow and Metabolism, 44(5), 660–679.

Kaplan, C. M., Kelleher, E., Irani, A., Schrepf, A., Clauw, D. J., & Harte, S. E. (2024). Deciphering nociplastic pain: clinical features, risk factors and potential mechanisms. Nature Reviews. Neurology, 20(6), 347–363.

Kosek, E., Clauw, D., Nijs, J., Baron, R., Gilron, I., Harris, R. E., Mico, J.-A., Rice, A. S. C., & Sterling, M. (2021). Chronic nociplastic pain affecting the musculoskeletal system: clinical criteria and grading system. Pain, 162(11), 2629–2634.

Mayer, T. G., Neblett, R., Cohen, H., Howard, K. J., Choi, Y. H., Williams, M. J., Perez, Y., & Gatchel, R. J. (2012). The development and psychometric validation of the central sensitization inventory. Pain Practice: The Official Journal of World Institute of Pain, 12(4), 276–285.

Murata, S., Hagiwara, A., Kaga, H., Someya, Y., Nemoto, K., Goto, M., Kamagata, K., Irie, R., Hori, M., Andica, C., Wada, A., Kumamaru, K. K., Shimoji, K., Otsuka, Y., Hoshito, H., Tamura, Y., Kawamori, R., Watada, H., & Aoki, S. (2022). Comparison of brain volume measurements made with 0.3- and 3-T MR imaging. Magnetic Resonance in Medical Sciences: MRMS: An Official Journal of Japan Society of Magnetic Resonance in Medicine, 21(3), 517–524.

Neblett, R., Sanabria-Mazo, J. P., Luciano, J. V., Mirčić, M., Čolović, P., Bojanić, M., Jeremić-Knežević, M., Aleksandrić, T., & Knežević, A. (2024). Is the Central Sensitization Inventory (CSI) associated with quantitative sensory testing (QST)? A systematic review and meta-analysis. Neuroscience and Biobehavioral Reviews, 161(105612), 105612.

Neumann, N., Domin, M., Schmidt, C.-O., & Lotze, M. (2023). Chronic pain is associated with less grey matter volume in the anterior cingulum, anterior and posterior insula and hippocampus across three different chronic pain conditions. *European Journal of Pain (London*, England), 27(10), 1239–1248.

Nishigami, T., Tanaka, K., Mibu, A., Manfuku, M., Yono, S., & Tanabe, A. (2018). Development and psychometric properties of short form of central sensitization inventory in participants with musculoskeletal pain: A cross-sectional study. PloS One, 13(7), e0200152.

Rouch, I., Edjolo, A., Laurent, B., Pongan, E., Dartigues, J.-F., & Amieva, H. (2021). Association between chronic pain and long-term cognitive decline in a population-based cohort of elderly participants. Pain, 162(2), 552–560.

Someya, Y., Tamura, Y., Kaga, H., Nojiri, S., Shimada, K., Daida, H., Ishijima, M., Kaneko, K., Aoki, S., Miida, T., Hirayama, S., Konishi, S., Hattori, N., Motoi, Y., Naito, H., Kawamori, R., & Watada, H. (2019). Skeletal muscle function and need for long-term care of urban elderly people in Japan (the Bunkyo Health Study): a prospective cohort study. BMJ Open, 9(9), e031584.

Treede, R.-D., Rief, W., Barke, A., Aziz, Q., Bennett, M. I., Benoliel, R., Cohen, M., Evers, S., Finnerup, N. B., First, M. B., Giamberardino, M. A., Kaasa, S., Korwisi, B., Kosek, E., Lavand’homme, P., Nicholas, M., Perrot, S., Scholz, J., Schug, S., … Wang, S.-J. (2019). Chronic pain as a symptom or a disease: the IASP Classification of Chronic Pain for the International Classification of Diseases (ICD-11). Pain, 160(1), 19–27.

Tzourio-Mazoyer, N., Landeau, B., Papathanassiou, D., Crivello, F., Etard, O., Delcroix, N., Mazoyer, B., & Joliot, M. (2002). Automated anatomical labeling of activations in SPM using a macroscopic anatomical parcellation of the MNI MRI single-subject brain. NeuroImage, 15(1), 273–289.

Uddin, L. Q. (2015). Salience processing and insular cortical function and dysfunction. Nature Reviews. Neuroscience, 16(1), 55–61.

Wang, J., Hill-Jarrett, T., Buto, P., Pederson, A., Sims, K. D., Zimmerman, S. C., DeVost, M. A., Ferguson, E., Lacar, B., Yang, Y., Choi, M., Caunca, M. R., La Joie, R., Chen, R., Glymour, M. M., & Ackley, S. F. (2024). Comparison of approaches to control for intracranial volume in research on the association of brain volumes with cognitive outcomes. Human Brain Mapping, 45(4), e26633.

Wardlaw, J. M., Smith, E. E., Biessels, G. J., Cordonnier, C., Fazekas, F., Frayne, R., Lindley, R. I., O’Brien, J. T., Barkhof, F., Benavente, O. R., Black, S. E., Brayne, C., Breteler, M., Chabriat, H., Decarli, C., de Leeuw, F.-E., Doubal, F., Duering, M., Fox, N. C., … STandards for ReportIng Vascular changes on nEuroimaging (STRIVE v1). (2013). Neuroimaging standards for research into small vessel disease and its contribution to ageing and neurodegeneration. Lancet Neurology, 12(8), 822–838.

Wolfe, F., Clauw, D. J., Fitzcharles, M.-A., Goldenberg, D. L., Häuser, W., Katz, R. L., Mease, P. J., Russell, A. S., Russell, I. J., & Walitt, B. (2016). 2016 Revisions to the 2010/2011 fibromyalgia diagnostic criteria. Seminars in Arthritis and Rheumatism, 46(3), 319–329.

Zhao, W., Zhao, L., Chang, X., Lu, X., & Tu, Y. (2023). Elevated dementia risk, cognitive decline, and hippocampal atrophy in multisite chronic pain. Proceedings of the National Academy of Sciences of the United States of America, 120(9), e2215192120.

